# Time-Dependent Image Restoration of Low-SNR Live Cell Ca^2+^ Fluorescence Microscopy Data

**DOI:** 10.1101/2021.10.05.462864

**Authors:** Lena-Marie Woelk, Sukanya A. Kannabiran, Valerie Brock, Christine E. Gee, Christian Lohr, Andreas H. Guse, Björn-Philipp Diercks, René Werner

## Abstract

Live cell Ca^2+^ fluorescence microscopy is a cornerstone of cellular signaling analysis and imaging. The demand for high spatial and temporal imaging resolution is, however, intrinsically linked to a low signal-to-noise ratio (SNR) of the acquired spatio-temporal image data, which impedes subsequent image analysis. Advanced deconvolution and image restoration algorithms can partly mitigate the corresponding problems, but are usually defined only for *static* images. Frame-by-frame application to spatio-temporal image data neglects inter-frame contextual relationships and temporal consistency of the imaged biological processes. Here, we propose a variational approach to *time-dependent* image restoration built on entropy-based regularization specifically suited to process low- and lowest-SNR fluorescence microscopy data. The advantage of the presented approach is demonstrated by means of four data sets: synthetic data for in-depth evaluation of the algorithm behavior; two data sets acquired for analysis of initial Ca^2+^ microdomains in T cells; and, to illustrate transferability of the methodical concept to different applications, one dataset depicting spontaneous Ca^2+^ signaling in jGCaMP7b-expressing astrocytes. To foster re-use and reproducibility, the source code is made publicly available.

## 1 Introduction

T cell activation represents the on-switch of the adaptive immune system [1]. Within tens of milliseconds after activation, highly dynamic, spatio-temporally restricted Ca^2+^ signals, termed Ca^2+^ microdomains, start occurring [1, 2], but the molecular machinery underlying this process still remains elusive. To better understand the principles of the formation and the temporal propagation of these signals as well as the contributions and roles of different components, high resolution live cell fluorescence microscopy is required, ideally implemented with both the spatial and the temporal resolution as high as possible. However, high resolution Ca^2+^ imaging has severe limitations: Low photon doses due to photo toxicity and photo bleaching as well as the per-se fugitive nature of Ca^2+^ signals in combination with out-of-focus light lead to an intrinsically low signal-to-noise ratio (SNR) [3]. This, in turn, significantly impedes identification and detailed analysis of Ca^2+^ microdomains and their spatio-temporal architecture.

The analysis of initial Ca^2+^ microdomains in T cells and the corresponding need to reliably identify related signaling events in live cell imaging data with poor SNR forms the basis and motivation of the present work, but it is only one example application. The general challenge to extract meaningful information from low SNR image time series data is applicable to many applications in the context of spatio-temporal fluoresence microscopy.

Techniques to increase the quality of captured images are typically referred to as image restoration or deconvolution.

In recent years, microscopy image restoration has been tackled increasingly using deep learning methods [4, 5], but a systematic problem with these approaches remains the risk of hallucinations, i.e. generation of structures not present in the acquired imaging data [6]. In addition, extensive amounts of suitable training data are usually required, limiting their applicability.

Conventional approaches, in contrast, work purely on the image data to be processed. They include classic, straight-forward methods such as nearest neighbor deconvolution or naive inverse filtering, which are computationally inexpensive but have drawbacks such as poor noise reduction and introduction of ringing artefacts. More sophisticated methods are often formulated as iterative algorithms and variational models, with a variety of data fidelity and regularization terms being proposed in literature. Most common approaches are (regularized) inverse filtering, including, e.g., Wiener filtering [7, 8], and (regularized) Lucy-Richardson (LR) deconvolution [9, 10]. For an overview please refer to, e.g., [11, 12].

Most functional formulations are, however, rather general, and the resulting algorithms perform poorly in low SNR scenarios [13]. In 2013, Arigovindan et al. introduced a functional formulation that was tailored to the specific characteristics of fluorescence microscopy [13]. In particular, they proposed an entropy-like formalism in combination with a second order derivatives-based regularization functional that suppresses noise but still preserves object details. A central rationale behind their formulation was, e.g., that, in contrast to general imaging data, in fluorescence microscopy data, ”high intensity points are more sparsely distributed and are co-localized with high-magnitude derivative points” [13]. The presented results were impressive especially for low SNR conditions. Yet, although motivated by demands of spatio-temporal imaging, the proposed formulation addressed only frame-by-frame deconvolution, i.e. the resulting algorithm was to be applied independently to each frame. While this is common to most image restoration methods (both deep learning and conventional approaches), recent work illustrated benefits of taking into account the spatio-*temporal* nature of the acquired data [14, 15].

In the present work, we extend the principle of entropy-like deconvolution proposed by [13] and suggest a novel variational approach tailored to image restoration of spatio-temporal fluoresence microscopy. The proposed approach utilizes the temporal information available in the imaging data to further improve image quality and SNR at low exposure times, thus enabling imaging with a higher temporal resolution. To foster re-use of the developed methods, the source code is freely available at `github/ipmi-icns-uke/XXX` (available after acceptance of manuscript). The repository also covers an implementation of the approach described in [13] to be applied to static microscopy data (no publicly available source code provided with the original publication). The advantage of the proposed approach is illustrated for four datasets. The first three datasets are related to the analysis of Ca^2+^ microdomains in T cells: (1) a synthetic data set with simulated Ca^2+^ signals and noise patterns to systematically evaluate the algorithm performance; (2) super-resolution spinning disc microscope image sequences acquired with a genetically encoded Ca^2+^ indicator tagged to a lysosomal channel in Jurkat T cells; and (3) widefield microscopy imaging of free cytosolic Ca^2+^ concentration ([Ca^2+^]_i_) in primary murine WT T cells. The fourth dataset illustrates the use in different application context, here confocal data of spontaneous Ca^2+^ signals in branches of an astrocyte in a mouse brain slice.

## 2 Materials and Methods

### 2.1 Mathematical Formulation

Following the concept of variational image deconvolution, the proposed approach builds on a common quadratic data fidelity term [13, 12], but extended to spatio-temporal image data, i.e. ||**Hv**_*t*_ − **w**_*t*_||^2^, where **w**_*t*_ is the measured image at time or frame *t*, **H** is the distortian matrix, and **v**_*t*_ the sought solution. Direct minimization of the data fidelity term would, in practice, suppress high frequency components of the solution and poses, mathematically, an ill-posed problem.

Thus, additional regularizing terms need to be included in the functional to be minimized. As described in the introduction, in [13], an algorithm specifically tailored to fluorescence microscopy was introduced. The main innovation was using second derivatives for a spatial smoothness enforcing regularization functional in an entropy-like term. Here, we expanded this approach to also include a functional term that enforces smoothness in the time domain. Similar to the motivation formulated for the spatial domain in [13] – sparsity of high intensity signals and high-magnitude derivatives – which we consider also applicable to the *temporal* characteristics of, e.g., Ca^2+^ microdomains, we also chose an entropy-like structure for the temporal regularization functional.

The overall minimization problem is given by

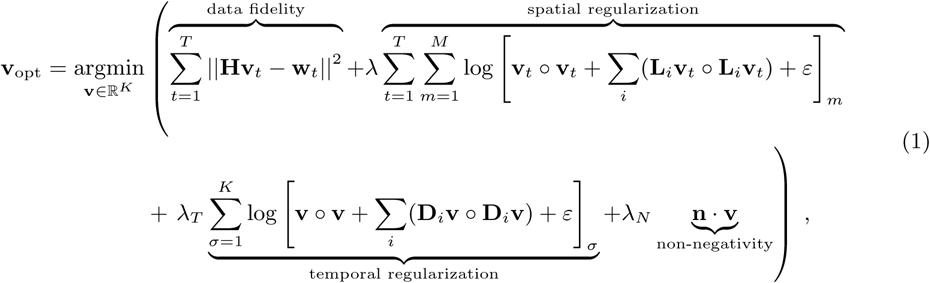

where **v** ∈ ℝ^*K*^ is a vectorized processed time series comprised of *T* frames, each a *N*_*x*_×*N*_*y*_ dimensional image, *M* = *N*_*x*_ · *N*_*y*_ and *K* = *M* · *T*. Each individual frame is denoted by the subscript *t* ∈ {1, …, *T*}, i.e. **v**_*t*_ ∈ ℝ^*M*^. The *measured* image time series is vectorized in the variable **w** ∈ ℝ^*K*^, with **w**_*t*_ ∈ ℝ^*M*^ being the vectorized time frame at time *t* and **v**_opt_ ∈ ℝ^*K*^ refers to the sought solution in terms of optimality with regard to the defined functional. The operator and distortian matrix **H** ∈ ℝ^*M*×*M*^ is in our case the point spread function (PSF) in Toeplitz form.

The operators **L**_*i*_ ∈ ℝ^*M*×*M*^ in the spatial regularization term represent the discretized second derivative filters in spatial directions, where the sum over *i* runs over *∂*^2^/*∂x*^2^, *∂*^2^/*∂y*^2^ and *∂*^2^/*∂x∂y*. The operator · ∘ · refers to the Hadamard, or element-wise, product. In the temporal regularization term, the **D**_*i*_ ∈ ℝ^*K*×*K*^ refer to the discretized second derivative filters containing the derivatives with respect to time. In this case, the sum over *i* runs over *∂*^2^/*∂t*^2^, *∂*^2^/*∂x∂t* and *∂*^2^/*∂y∂t*.

The sum over *m* runs over all pixel within one time frame, so *m* ∈ {1…*M* ≡ *N*_*x*_ · *N*_*y*_}, while *σ* is a composite index referring to a pixel within a specific time frame and running over all pixels in all time frames, i.e. *σ* ∈ {1…*K* ≡ *N*_*x*_ · *N*_*y*_ · *T*}. In our notation, the value of a pixel in a specific time frame can thus be addressed by either *v*_*σ*_ or [**v**_*t*_]_*m*_.

The vector **n** ∈ ℝ^*K*^ ensures positivity of the result and contains the entries *n*_*σ*_ where *n*_*σ*_ = 0 if *v*_*σ*_ ≥ 0 and 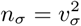 if *v*_*σ*_ < 0.

The parameters *λ, λ*_*T*_ and *λ*_*N*_ are Lagrange parameters to weight the regularization terms. They are to be determined empirically. *ε* is a small positive constant to avoid occurrence of log (0) in the regularization terms.

The first term in the cost function, i.e. the data fidelity term in eq. (1), ensures the agreement with the forward model, i.e. the image distortion **w**_*t*_ = **Hv**_*t*_. The second term, controlled with Lagrange parameter *λ* and denoted as spatial regularization in eq. (1), is known from [13] and denotes the regularization functional enforcing smoothness within the spatial dimensions of the image. New in our proposed method is the third term proportional to *λ*_*T*_. This regularization functional enforces smoothness *over time*. The last term proportional to *λ*_*N*_ is a standard term to avoid negative pixel values in the resulting image.

The optimal solution of problem eq. (1) is found using an iterative minimization algorithm detailed in appendix A.

Note that the *static* entropy-based image restoration algorithm described in [13] is also included as a limit for *λ*_*T*_ → 0 in the above description. Whenever this algorithm is referred to in the following for comparison purposes, this means our implementation with *λ*_*T*_ set to zero. We choose as abbreviations *ER* for the static entropy deconvolution and *TD ER* for the time-dependent entropy deconvolution.

### 2.2 Experiments: Imaging Data and Evaluation

The performance of the proposed spatio-temporal deconvolution was tested and compared to static entropy-based deconvolution and standard LR deconvolution (implementation of the MATLAB Image Processing Toolbox 2019a) by means of four different datasets: a synthetic image data set, two fluorescence microscopy image datasets acquired in the context of Ca^2+^ microdomain analysis in T cells, and a last dataset acquired by confocal fluorescence Ca^2+^ imaging of an astrocyte in an acute mouse brain slice to illustrate transferability of the proposed methodical developments to a different application context.

#### 2.2.1 Data Set 1: Synthetic Image Data

Simulation of Ca^2+^ fluorescence microscopy time series data started on a black canvas. To generate a texture, Perlin noise was added [4]. The texture was used to place “glowing” spots (small Gaussian intensity peaks) in a randomly clustered manner. Afterwards, the Perlin noise was removed and the spots were moved over time according to Brownian motion.

To degrade the images for deconvolution evaluation purposes, they were first convolved with a PSF. The applied PSF was the same as used for the real microscopic data of dataset 3 (see below). Then, Poisson and Gaussian noise were added. The noise levels were varied to analyze the performance of the image restoration algorithms as a function of input image data SNR. This can also be interpreted to simulate different exposure times.

Poisson noise was varied by dividing the signal by a parameter before calculating the Poisson distribution. The result was re-scaled by this same parameter to preserve the original dynamic range. The Gaussian noise was varied by adding Gaussian noise with different variance. Since the exact values of these parameters are rather arbitrary in synthetic images, we scaled them to dimensionless noise levels to better illustrate the amount of noise present in the images. The exact parameters and generation methods can be seen in the published source code.

Different to subsequent real live cell microscopy imaging data, the synthetic imaging data allow for quantitative comparison of sought optimal images, i.e., the original input images before degradation by the PSF and noise application. For evaluation purposes, we calculated the structural similarity index (SSIM) between patches of the original images **v**_orig_ and the restored images **v**_dec_, given by

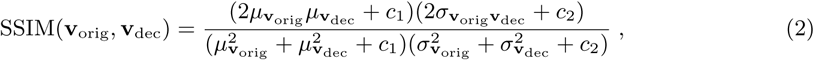

where *µ*_**x**_ denotes the average and 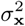 the variance of the intensity values of image patch **x**, while *σ*_**xy**_ denotes the covariance between two image patches **x** and **y**. *c*_1_, *c*_2_ are small constants.

Moreover, as an approach that requires no ground truth reference image for quantitative assessment of image restoration success, the Gaussian noise variance of the image background was estimated according to the patch-based approach presented in [16].

#### 2.2.2 Data Set 2: Genetically encoded Ca^2+^ indicator for optimal imaging (GECO) tagged to lysosomal TPC2 in Jurkat T cells

The second data set was acquired in the context of the analysis of the role of Ca^2+^ release processes during the formation of initial Ca^2+^ microdomains in T cells. Jurkat T cells were transiently transfected with two pore channel 2 (TPC2) fused to a low affinity genetically encoded Ca^2+^ indicator for optical imaging (GECO-1.2). Previously, this GECO was tagged to ORA1 channels in the plasma membrane and only Ca^2+^ entry through Orai1 was visualized [17]. Here, only Ca^2+^ release from the lysosomes through TPC2 should be detected.

The images were acquired with a 100-fold magnification objective (Zeiss) fitted in a super-resolution spinning disc microscope (Visitron) and a scientific complementary metal-oxide-semiconductor camera (Orca-Flash 4.0, C13440-20CU, Hamamatsu). Different times of acquisition were used for time lapse series (100 ms, 150 ms, 200 ms, 400 ms), a 561 nm laser adopted to excite TPC2-R.GECO.1.2, and the emission wavelength was detected at 606/54 nm.

#### 2.2.3 Data Set 3: Free cytosolic Ca^2+^ concentration imaging in primary murine WT T cells

The third data set depicts the free cytosolic Ca^2+^ concentration ([Ca^2+^]_i_) of primary murine WT T cells before and immediately after T cell activation. Imaging was performed as detailed in [1, 2]. The cells were loaded with Fluo4-AM and Fura Red-AM. For T cell stimulation, protein G beads (Merck Millipore) were coated with antibodies (anti CD3/anti CD28). Image sequences were acquired on a Leica IRBE2 widefield microscope using 100-fold magnification. A Sutter DG-4 was used as light source, and the image sequence frames acquired with an electron multiplying charge-coupled device camera (C9100, Hamamatsu). A dual-view module (Optical Insights, PerkinElmer Inc.) was used to split the emission wavelengths (filters: excitation 480/40; beam splitter, 495; emission 1, 542/50; emission 2, 650/57). Exposure time was 25 ms (image acquisition with 40 frames/s).

#### 2.2.4 Data Set 4: Confocal Ca^2+^ imaging in astrocytes *in situ*

The fourth data set was aquired in an astrocyte in an acute mouse brain slice. The genetically encoded Ca^2+^ indicator jGCaMP7b (Addgene # 171118) was subcloned into a AAV-PhP.eB vector under control of the GFAP promoter and viruses were systemically applied by retrobulbar injection [18]. After 14 days, jGCaMP7b-expressing astrocytes were visualized with a confocal fluorescence microscope (eC1, Nikon, Düsseldorf, Germany; equipped with a 16x objective, NA 0.8), in acute brain slices of the olfactory bulb using a 488 nm laser for excitation (emission filter 515/15) at a frame rate of 1 Hz.

## 3 Results

### 3.1 Synthetic Data

A comparison of the effects of the different deconvolution approaches for an example of the generated synthetic data with different noise levels is given in fig. 1. The simulated ground truth data are shown in fig. 1(**a**).

**Figure 1:**
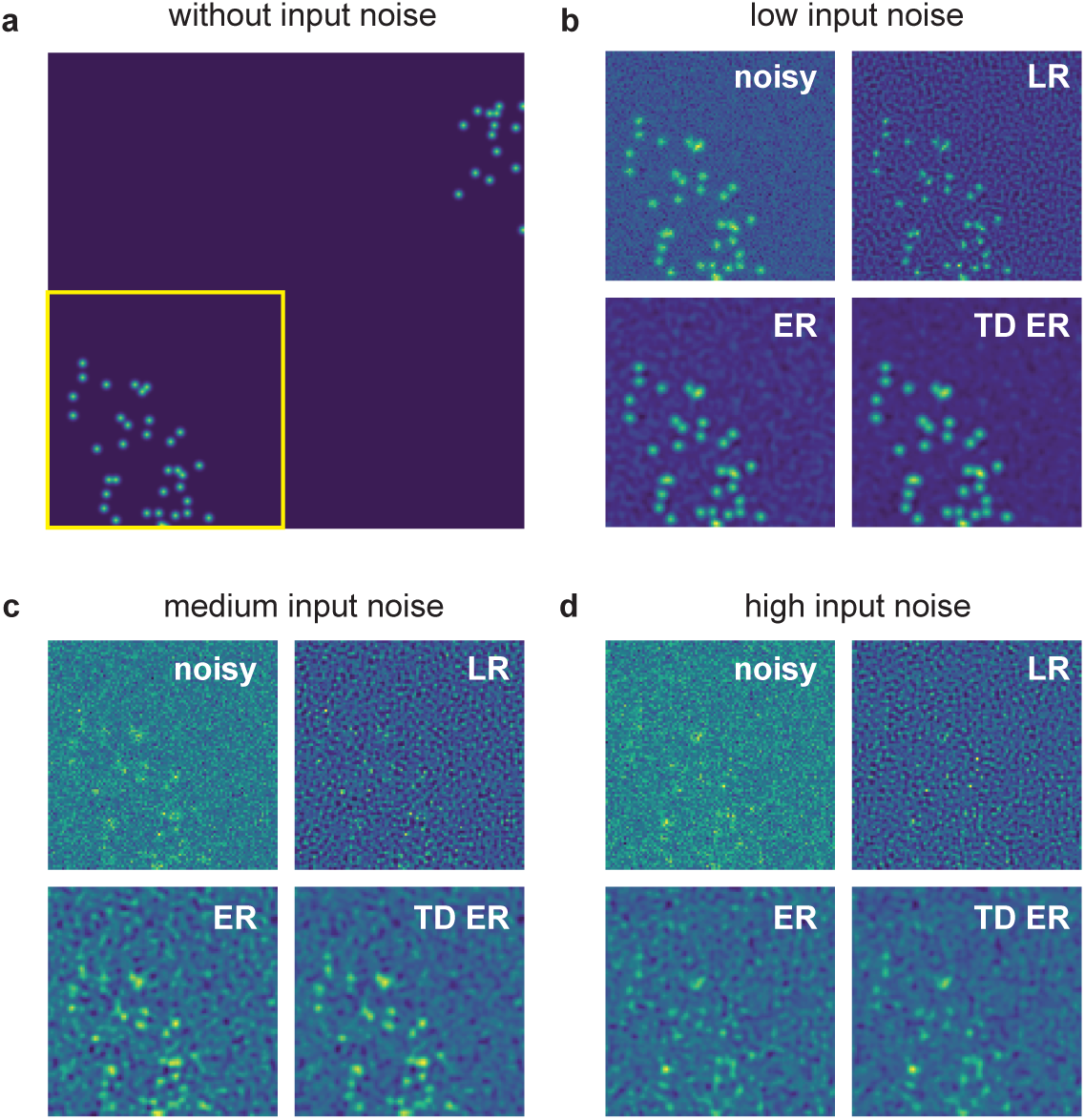
Example of synthetic data processed with the different deconvolution methods. (**a**) Sample frame of synthetic data without any added noise and before applying the PSF. The yellow box indicates the region of interest pictured in panels (**b**)-(**d**), which show input noisy images for various noise levels as well as image restoration results. The parameters for the entropy deconvolution are *λ* = 0.1 and (TD ER: *λ* = 0.1, *λ*_*T*_ = 0.1) and *ε* = 0.001. LR: Lucy Richardson deconvolution; ER: entropy regularization-based deconvolution (static); TD ER: time-dependent entropy regularization-based deconvolution.

The region of interest (ROI) indicated by the yellow box is then focused on in the panels (**b**)-(**d**), which are all structured in the same way: The left upper image represents the noisy ROI of the image that is input into the deconvolution algorithms. The other images are the corresponding restored image ROI for LR deconvolution (right upper image), static entropy-based deconvolution (ER, left lower image), and the proposed temporal entropy-based deconvolution (TD ER, right lower image). The input noise levels are as follows: (**b**) low noise, (**c**) medium noise and (**d**) high noise.

For all noise levels, the proposed time-dependent algorithm presents the least amount of background noise and highest SNR after image restoration, with the discrepancy between time-dependent and static deconvolution becoming most evident for high noise level (i.e., low-SNR) input data as in panel (**d**). In contrast to the entropy-based approaches, LR deconvolution tends to introduce ringing artefacts to the result, which amplify the background noise for low-SNR input data. Merely for the lowest noise level shown in panel (**b**), the LR algorithm performs best, as it sharpens the signal while the entropy algorithms tend to blur it instead.

The corresponding quantitative analysis is presented in fig. 2, showing the mean normalized SSIM as well as the estimated background noise for the image restoration approaches as a function of the input Gaussian noise level and averaged over different Poisson noise levels.

**Figure 2:**
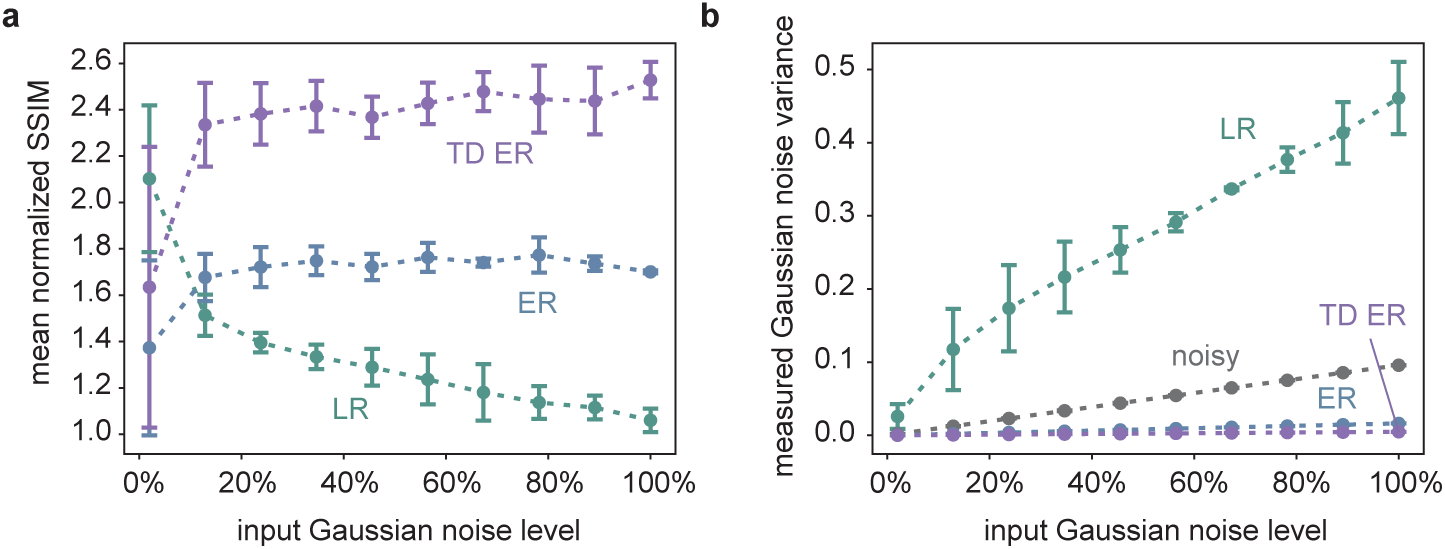
(**a**) shows the SSIM of the different image restoration methods, averaged over all time frames of ten different, randomly generated synthetic datasets like the one in fig. 1(**a**), normalized with respect to the SSIM between the noisy and the original image as a function of the input Gaussian noise. Thus, a value larger than one indicates image quality improvement compared to the noisy input image. The measurement points correspond to average values obtained for three different Poisson noise levels, and the error bars indicate the influence of varying the Poisson noise levels in terms of the standard deviation of the respective different simulations. (**b**) represents the results of the Gaussian noise estimation as a function of the input Gaussian noise for the different deconvolution methods as well as for the original noisy image, where the latter is pictured in greys and represents a plausibility check of the applied noise estimation approach.

The SSIM is calculated according to eq. (2) for the individual frames of the restored images and the original input data and averaged over all time frames. In the diagram, respective SSIM values are normalized by dividing the SSIM obtained for an image restoration approach by the SSIM for comparison of the noisy input data and the underlying original data. Thus, SSIM values larger than one indicate an image improvement in terms of SSIM.

The quantitative data support the visual impression. The TD ER algorithm performs best, except for very low noise values, where the LR deconvolution reveals higher SSIM values. With increasing noise, image quality improvement by LR deconvolution drastically decreases in terms of SSIM (normalized SSIM values ≈ 1; a value of 1 indicates similar SSIM of the noisy input image and the restored image). Better results are obtained using the entropy approaches (ER: normalized SSIM of approximately 1.7; TD ER: normalized SSIM of approximately 2.4), which are optimized for processing low SNR fluorescence microscopy data. The amount of Poisson noise has a comparatively small influence on the result, as can be seen by the error bars in eq. (2)(**a**), which show the standard deviation for processing similar image series with different Poisson noise levels.

The estimated Gaussian noise variance of the image background depicted in fig. 2(**b**) is in line with the SSIM results and the visual impression.

The grey line represents a consistency check of the automated background noise estimation method, as it shows the estimated Gaussian noise variance of the noisy input image as a function of the input Gaussian noise variance. The linear relationship indicates reliability of respective results.

Both the ER and the TD ER algorithms considerably decrease the measured background noise, while the LR algorithm appears to magnify it with increasing input noise level, in agreement with the visual impression of fig. 1(**b**)-(**c**). The LR results are also the only ones that depended on the input Poisson noise level, with the standard deviation indicating this influence as explained and visualized in fig. 2(**a**). The remaining background noise of the images obtained by both entropy algorithms as well as the background noise of the original noisy images differ only little for different Poisson levels, and respective error bars are too small to be pictured in fig. 2(**a**). Overall, the least amount of background noise is present in the images generated using the TD ER algorithm.

### 3.2 Live cell fluorescence microscopy

In fig. 3, representative frames of the acquired image sequences of data set 2 and corresponding outputs of the deconvolution approaches are shown. The data represent Ca^2+^ release through the two pore channel 2 (TPC2) on lysosomes in Jurkat T cells using TPC2-R.GECO.1.2. With this low affinity genetically encoded Ca^2+^ indicator, only Ca^2+^ at the mouth of the TPC pore can be visualized.

**Figure 3:**
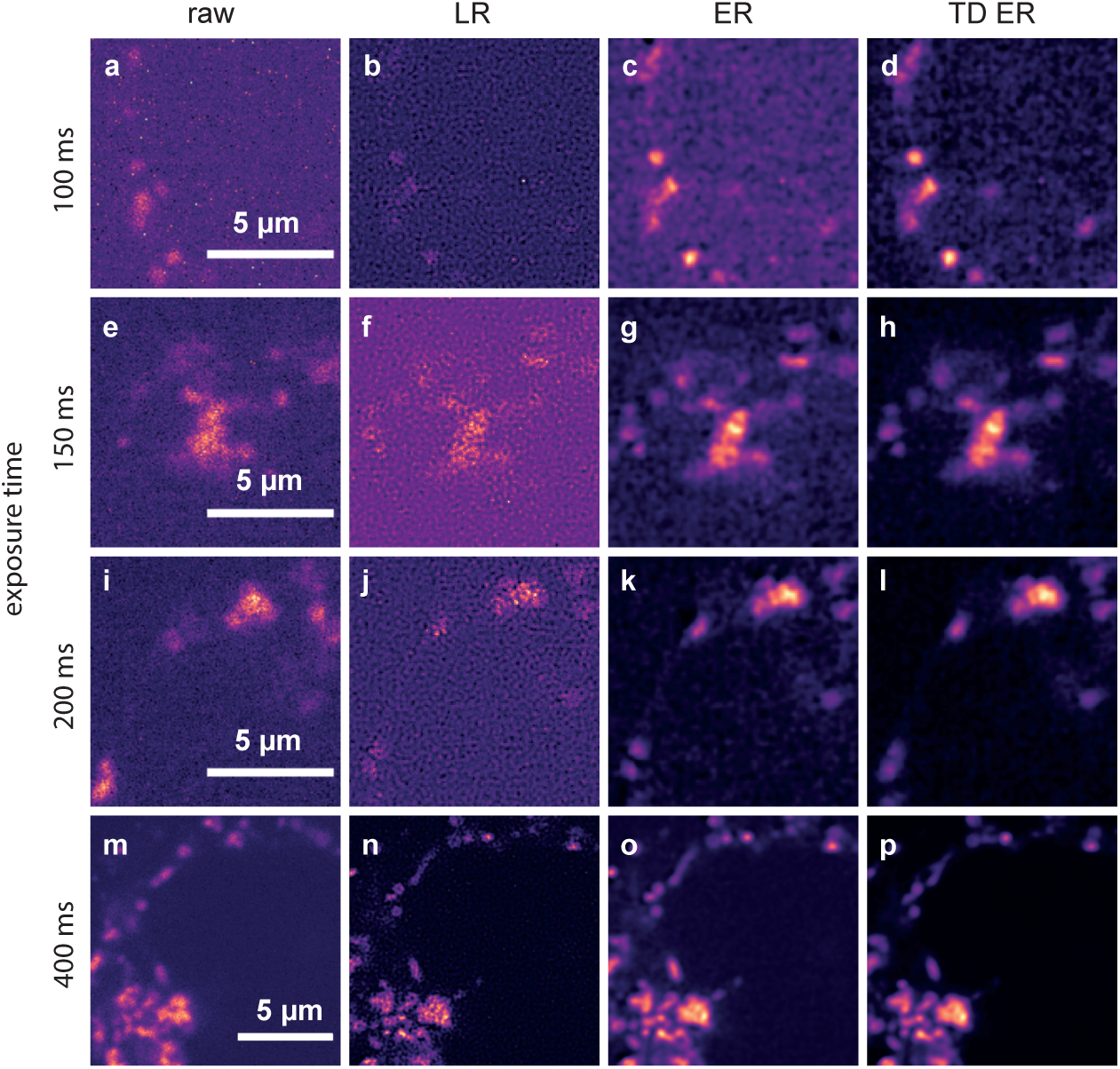
Comparison of the different deconvolution methods for TPC2-R.GECO.1.2 images captured with different exposure times. (**a**), (**e**), (**i**), (**m**): raw data, captured at 100 ms, 150 ms, 200 ms, and 400 ms; (**b**), (**f**), (**j**), (**n**): images deconvolved using MATLAB’s Lucy-Richardson (LR) algorithm; (**c**), (**g**), (**k**), (**o**): images deconvolved using the static entropy algorithm (ER); (**d**), (**h**), (**l**), (**k**): images deconvolved with the time-dependent entropy algorithm (TD ER). Parameters for the entropy algorithms are *λ* = 2.0 and (TD ER: *λ* = 2.0, *λ*_*T*_ = 2.0) and *ε* = 0.001.

The first column shows raw images captured with exposure times of 100 ms (**a**), 150 ms (**e**), 200 ms (**i**) and 400 ms (**m**). The second column depicts corresponding deconvolution results obtained with the MATLAB implementation of the LR algorithm for these exposure times. Images restored by the ER deconvolution (*λ* = 2.0) are shown in the third column, and corresponding results for the proposed TD ER approach (*λ* = 2.0, *λ*_*T*_ = 2.0) are given in the fourth column. In both the static and time-dependent entropy algorithms, *ε* = 0.001.

It can be clearly seen that the noise level in the raw images increases significantly when reducing the exposure time from 400 ms to 100 ms. The Lucy Richardson deconvolution increases the noise level even further for lower exposure times, while the entropy-based algorithms perform much better (third and fourth column). The time-dependent algorithm (rightmost column) shows the least amount of noise while recovering much of the original signal. This effect is especially pronounced for the lower exposure times, where in the raw image, hardly any signal can be discerned, while our algorithm manages to recover a relatively clear signal. For very high exposure times, such as 400 ms shown in the last row of fig. 3, the improvement is, however, minimal at best. While even here background noise is reduced, the signal also appears slightly blurred.

A more detailed look at the signal recovery is given in fig. 4. Here, results for the different deconvolution methods are shown for a ROI of a time series captured with 100 ms exposure time. The first column, fig. 4(**a**), shows the raw image of a single time frame in total (first row) with a zoomed-in ROI below. Panel (**e**) represents an intensity profile along the blue line in the zoom plot. The same structure applies to the other columns, with (**b**), (**c**) and (**d**) showing the deconvolution results of the LR, ER and TD ER algorithms, respectively, and (**f**)-(**h**) the corresponding intensity profiles along the pictured blue lines. The intensity plots illustrate that the LR deconvolution seems to sharpen the signal, but also amplifies the noise; based on the intensity profile, it is difficult to distinguish signal from noise. The ER algorithm clearly recovers the signal while reducing the noise, and the TD ER further enhances the SNR.

**Figure 4:**
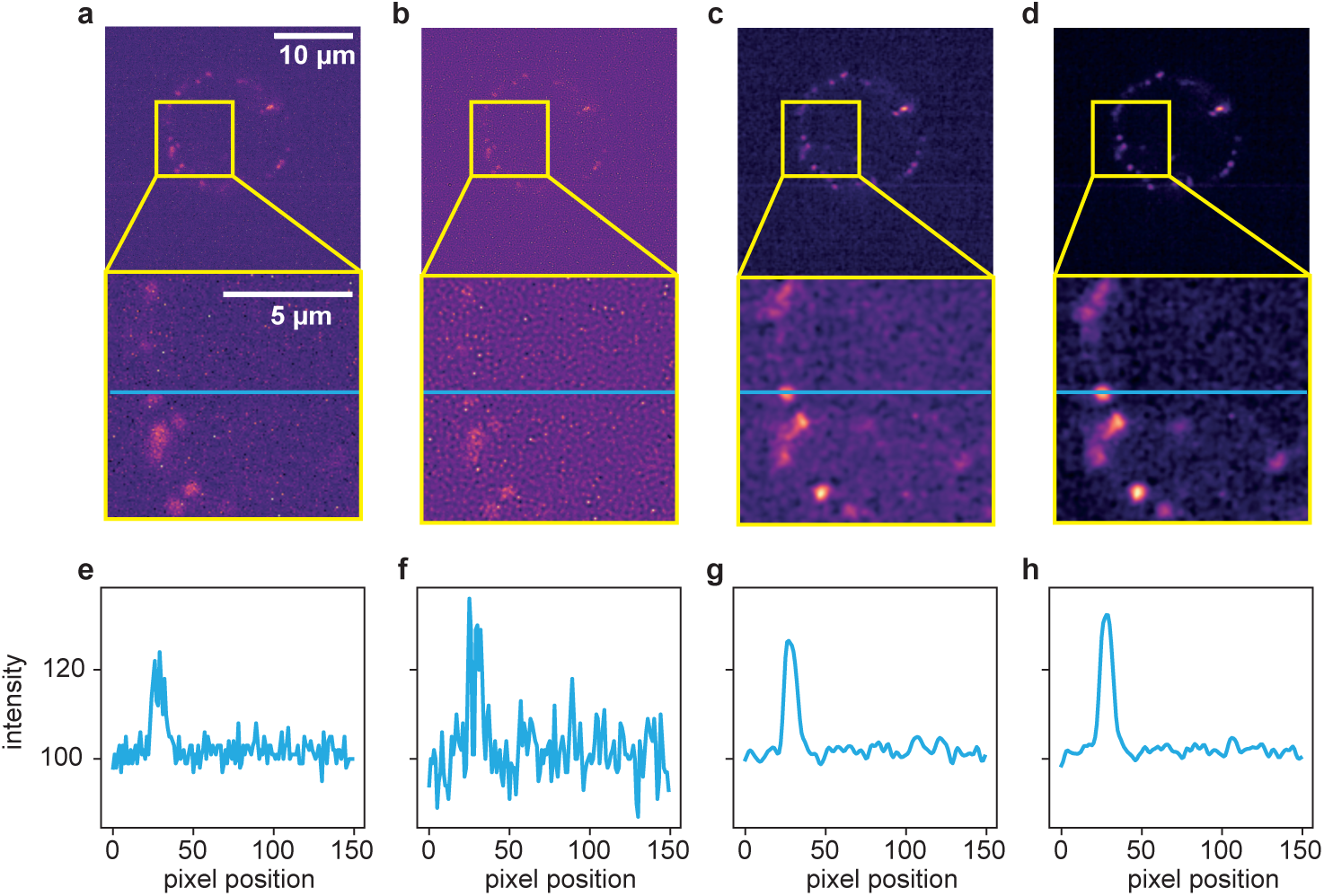
Comparison of the different deconvolution methods for a time series of data set 2, captured at 100 ms exposure time. (**a**) Raw image. (**b**) Deconvolved with the MATLAB Lucy-Richardson algorithm. (**c**) Deconvolved by ER with *λ* = 2.0. (**d**) Deconvolved with the proposed TD ER with (*λ* = 2.0, *λ*_*T*_ = 2.0). Each panel includes a zoomed-in region of interest indicated in yellow. (**e**), (**f**), (**g**), and (**h**) show the intensity profile plotted along the blue line in the frames above. All entropy-based algorithms here use *ε* = 0.001.

To further illustrate the potential of the proposed approach, two additional live cell imaging data sets were processed and analyzed.

Results obtained on Data set 3 are illustrated in fig. 5 and fig. 6. Shown in fig. 5(**a**) is an image frame captured using Fluo-4 as the indicator dye, while (**c**) depicts the simultaneously acquired image frame captured using Fura Red as the indicator dye. The frame corresponds to a time point shortly after T cell activation.

**Figure 5:**
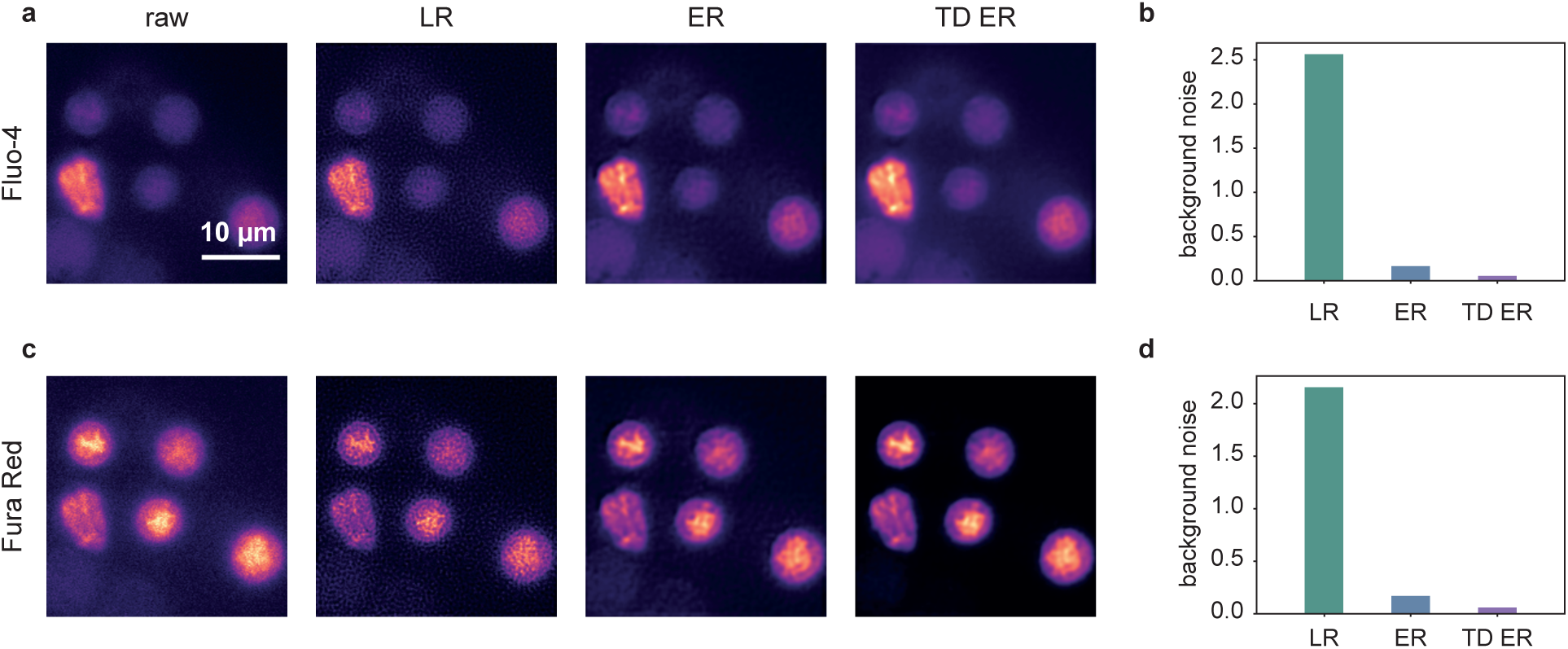
Deconvolution results for [Ca^2+^]_i_ imaging and frames (**a**) using Fluo-4 and (**c**) using FuraRed as the indicator dye. From left to right: raw image, LR result, ER result, and TD ER result. Entropy-based deconvolution parameters were *λ* = 0.4 (TD ER: *λ* = 0.4, *λ*_*T*_ = 0.4) and *ε* = 0.001. Panels (**b**) and (**d**) show the corresponding estimated background noise remaining in the deconvolved images, normalized to the background noise of the raw image for the different deconvolution methods.

**Figure 6:**
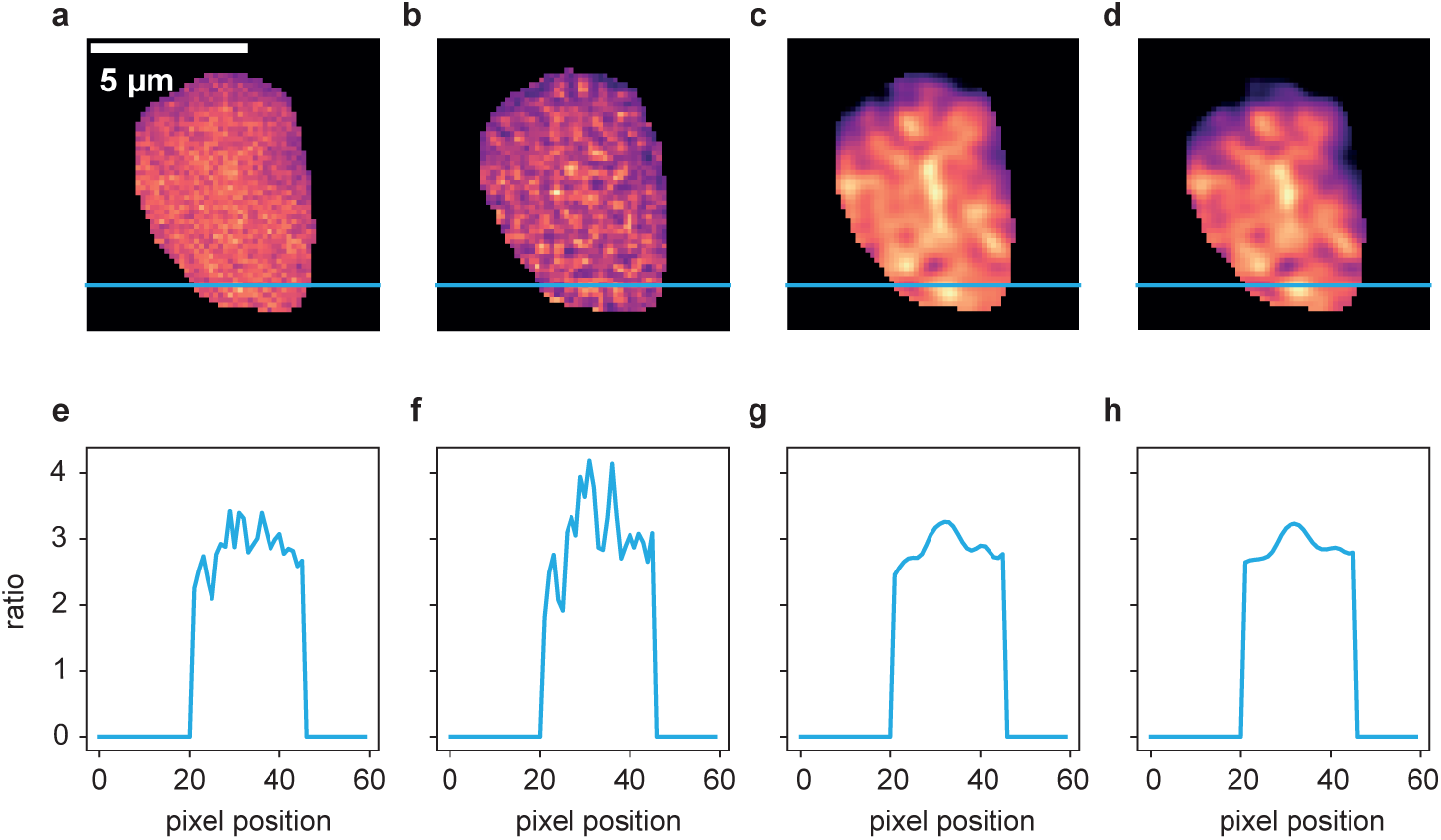
Ratio of the deconvolution results of the two channels from fig. 5 after postprocessing according to [2]. (**a**) Fluo-4/FuraRed ratio of raw images, (**b**) ratio of LR results, (**c**) ratio of ER results and (**d**) ratio of TD ER results. Panels (**e**) - (**h**) show the intensity profile plotted along the blue line in the frames above.

The panels show, from left to right, the original raw data and the deconvolution results obtained with the LR, the ER and the TD ER algorithms. Similar to above experiments and data sets, the least amount of background noise is present both visually in (**a**) and (**c**) and, in terms of estimated Gaussian background noise, quantitatively in (**b**) and (**d**) in the output images of the TD ER algorithm. In fact, entropy deconvolution eliminates the background noise almost entirely. The numbers given in (**b**) and (**d**) are shown as a ratio, i.e. the noise level after image restoration divided by the estimated noise level of the raw data. Thus, values < 1 represent a decrease of background noise.

Performing the postprocessing process as described in [2] (rigid registration of the two channel time series data, bleaching correction, cell segmentation), the two channel image data were then combined to ratio images representing the free cytosolic Ca^2+^ concentration, [Ca^2+^]_i_. One exemplary cell is shown in fig. 6, where (**a**)-(**d**) show the ratio images computed based on the aligned and processed raw images, the images after LR deconvolution, and after ER and TD ER image restoration. Panels (**e**)-(**h**) show the intensity profile along the blue line in the images above. While the ratio of the raw channels appears to be very grainy, the Ca^2+^ microdomains in the deconvolved images, in particular for ER and TD ER, can be much more easily distinguished from noise.

The results for data set 4, a jGCaMP7b-expressing astrocyte in a mouse brain slice, are shown in fig. 7, illustrating transferability of the developed approach to different application contexts than Ca^2+^ microdomain analysis. While the original SNR for the input data appears to be already quite good for the large and brightly labeled cell body, the fine cell branches barely stand out against the background. Here, the entropy algorithms both considerably decrease the amount of background noise, making it easier to separate the delicate structures of the astrocyte branches from the background. For a visual impression, see fig. 7(**a**)-(**d**) and supplementary videos S1-S4. Quantitatively, the background noise reduction is shown in fig. 7(**i**) in the measurement of the background noise variance of the deconvolved images, again normalized onto the original noise level. The intensity profile plotted in panels (**e**)-(**f**) further confirms that the SNR, while already acceptable in the raw image, is further improved by the entropy algorithms as shown in (**g**) and (**h**).

**Figure 7:**
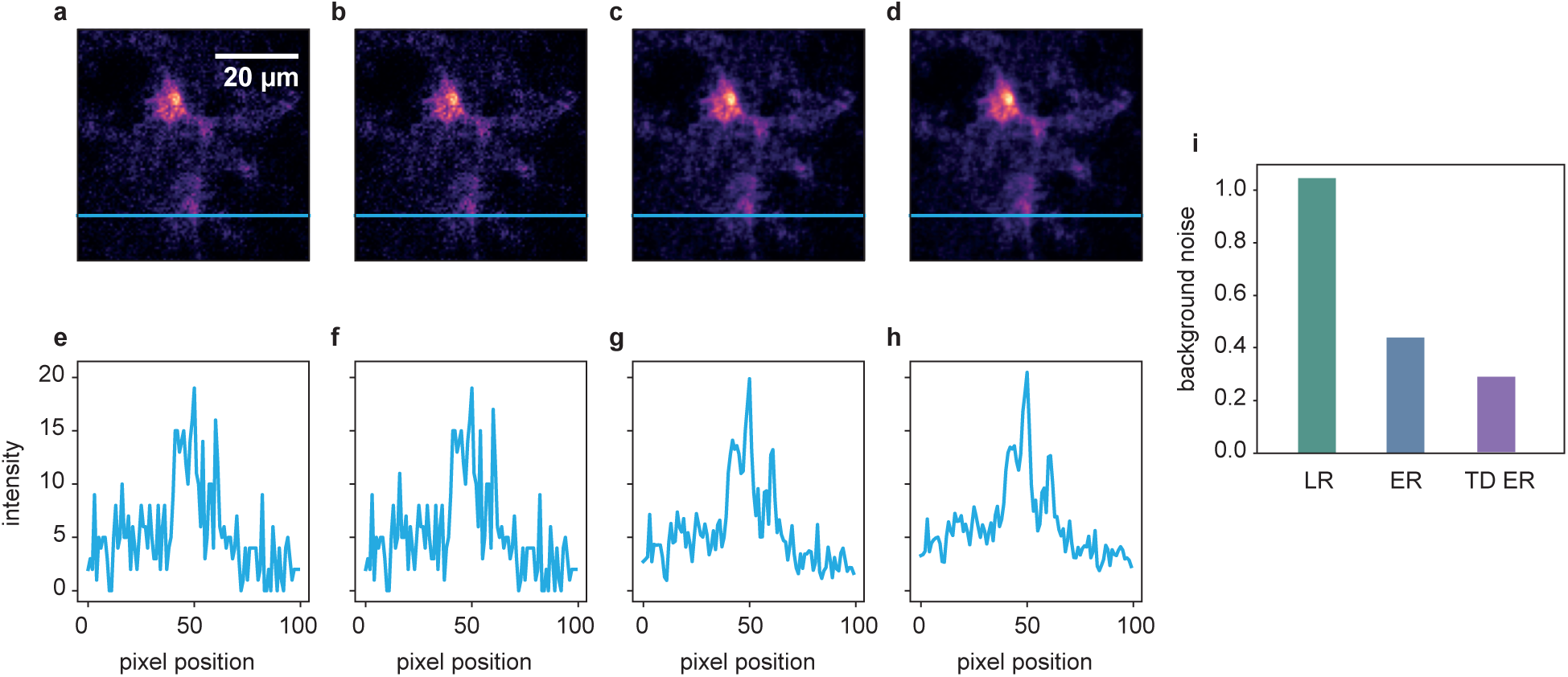
Deconvolution results for a jGCaMP7b-expressing astrocyte in a mouse brain slice. (**a**) raw image, (**b**) LR result, (**c**) ER result, and (**d**) TD ER result. Entropy parameters here are *λ* = 0.05 and (*λ* = 0.05, *λ*_*T*_ = 0.05) and *ε* = 0.001. Panels (**e**) - (**h**) show the intensity profile plotted along the blue line in the frames above. Panel (**i**) shows the amount of background noise remaining in the image after the application of the different deconvolution algorithms, normalized to the noise level of the original data.

The computation time for a 500×500 pixel time series with 100 frames using the two entropy algorithms ranges between a few seconds and a few minutes on a standard desktop PC, depending on the convergence of the algorithm, which in turn depends on the input data.

## 4 Discussion

Motivated by the intrinsically low SNR for live cell Ca^2+^ image sequences acquired by fluorescence microscopy at both high spatial resolution and high temporal resolution, we proposed integration of the temporal dimension of respective image data into variational image restoration. Method development built on an image restoration specifically tailored to particularities of fluorescence microscopy [13]. Here, (1) we extended the underlying entropy-based regularization and the dedicated numerical solving scheme to spatio-temporal image sequences, (2) demonstrated the superiority of the proposed time-dependent image restoration approach compared to static entropy-based image restoration and common LR deconvolution, and (3) made the source code publicly available.

Demonstration of the advantages of the proposed image restoration approach was based on synthetic as well as real live-cell Ca^2+^ imaging data, with two of the latter being acquired in the context of Ca^2+^ microdomain formation analysis after T cell activation, and one additional data set showing a jGCaMP7b-expressing astrocyte in a mouse brain slice. For all data sets, the observed effects were consistent: The time-dependent deconvolution considerably reduced the level of noise, in particular for low-SNR input image sequences. We therefore expect the approach to be promising for live cell imaging data acquired in different application context.

For high(er) SNR input image sequences, the quantitative evaluation has, however, shown that the performance of the common LR deconvolution is on par with both entropy-based image restoration approaches. Moreover, visually, the entropy approaches tend to blur spots of high Ca^2+^ concentration (also particularly visible for high-SNR input data). We hypothesize that this due to the present data fidelity term of the functional in eq. 1, and will in the future adjust the functional by changing the term from least squares to a Poisson noise-specific term, as low photon rates typical for fluorescence microscopy tend to obey Poisson statistics. This, however, requires a different numerical scheme and algorithm for the minimization of the overall functional and is beyond the scope of the present paper.

We encourage other researchers to test the proposed algorithm on their data and to contact us in case of problems.

## Supplementary Materials

The following supplementary materials are available online after acceptance: Videos S1-S4: movies corresponding to the astrocyte data shown in fig. 7; S1: raw data; S2: LR deconvolution; S3: ER deconvolution; S4: TD ER deconvolution.

## Author Contributions

Conceptualization, L.-M.W., A.H.G., B.-P.D., and R.W.; methodology, L.-M.W. and R.W.; software, L.-M.W.; validation, L.-M.W. and R.W.; formal analysis, L.-M.W.; investigation, L.-M.W., S.K., V.B., C.L., and R.W.; resources, all authors; data curation, L.-M.W.; writing—original draft preparation, L.-M.W., B.-P.D., C.L., and R.W.; writing—review and editing, all authors; visualization, L.-M.W.; supervision, R.W.; project administration, B.-P.D. and R.W.; funding acquisition, B.-P.D., R.W., A.H.G., C.E.G., and C.L. All authors have read and agreed to the published version of the manuscript.

## Funding

This research was funded by Deutsche Forschungsgemeinschaft (DFG, German Research Foundation) grant number 335447717 - SFB 1328, projects A01 to A.H.G., A02 to B.-P.D. and R.W., and A07 to C.E.G. and C.L.

## Data Availability Statement

The source code of proposed method and the synthetic data sets are publicly available at `github/ipmi-icns-uke/XXX` (available after acceptance of manuscript).

## Conflicts of Interest

The authors declare no conflict of interest. The funders had no role in the design of the study; in the collection, analyses, or interpretation of data; in the writing of the manuscript, or in the decision to publish the results.

## Abbreviations

ER: static entropy deconvolution
GECO: genetically encoded Ca^2+^ indicator for optimal imaging
LR: Lucy-Richardson
PSF: Point spread function
ROI: Region of interest
SNR: Signal-to-noise ratio
SSIM: Structural similarity index
TD ER: time-dependent entropy deconvolution
TPC: two pore channel

## A Algorithm Details

### A.1 Minimization of Cost Functional (1)

As described in the main text, our algorithm is conceptually an extension of the entropy deconvolution proposed by Arigovindan et al. [13]. There was, however, no corresponding source code available. We, therefore, developed the program completely from scratch with the ideas in [13] as a starting point. Introduced in section 2, the cost functional to minimize is given by

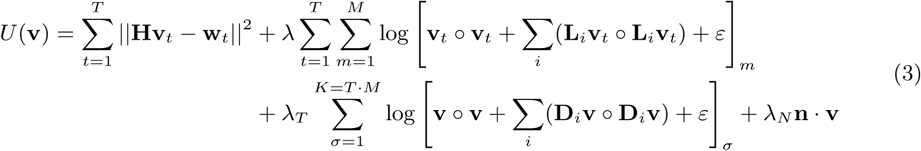

Taking the derivative and setting it to zero leads to the following minimality condition

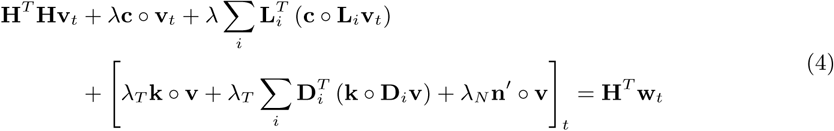

for the *t*-th time frame. The elements of the vector **c** are given by

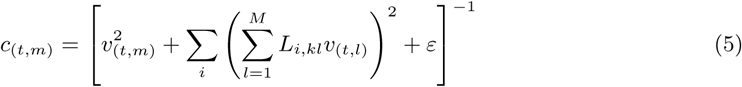

where (*t, m*) denotes a composite index to reference both the time frame *t* and pixel location *m*. The elements of the vector **k** are given by

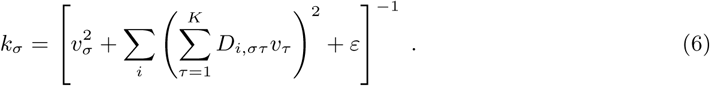

Since eq. (4) is solved iteratively, a starting condition is needed. It is useful to define an approximation which can be easily inverted. Extending the ansatz in [13], we choose the following initial condition for the *t*-th time frame

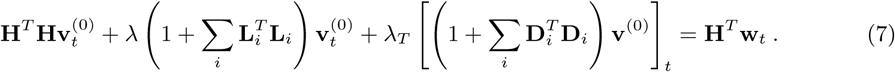

Note that the matrices **H, L**_*i*_, **D**_*i*_ are circulant to represent the convolution. Therefore they are diagonalized by the discrete Fourier transform. This leads to the following solution to eq. (7):

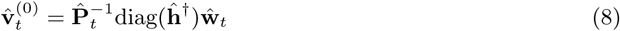

where

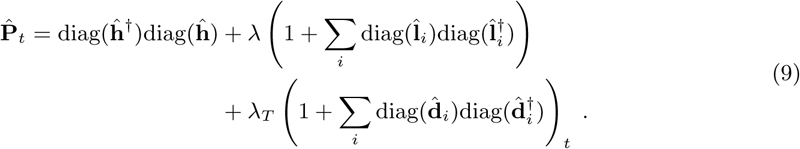

Here, 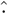 denotes the Fourier transform and diag(**x**) the diagonal matrix with entries **x** along the diagonal. For circulant matrices, the eigenvalues are given by one of its rows (all other rows are permutations) and the Fourier transform is simply a diagonal matrix with these values along the diagonal.

Going back to the full problem, the left hand side of eq. (4) can be re-written as a single operator as

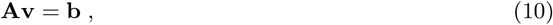

where **b**_*t*_ = **H**^*T*^ **w**_*t*_ and **v** denotes again the entire time series. The solution, i.e. **v**, can be found iteratively following the ansatz of [13] with

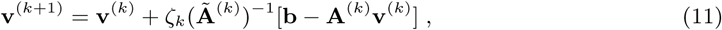

where *k* is the iteration index, *ζ*_*k*_ a damping factor, and **Ã** ^(*k*)^ an approximation of **A**^(*k*)^ that can be inverted easily. Here

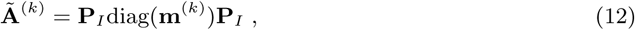

where **P**_*I*_ is the inverse Fourier transform of 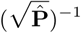, where 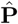 is given by eq. (9) and diag(**m**^(*k*)^) is the diagonal approximation of **A**^(*k*)^ with

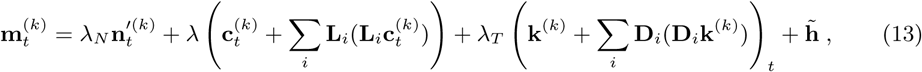

where the vectors **c, k** are given by eqs. (5) and (6) and the vector 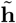 is constant with elements

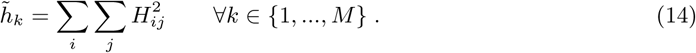

To facilitate notation in the algorithm, some abbreviations are introduced as follows

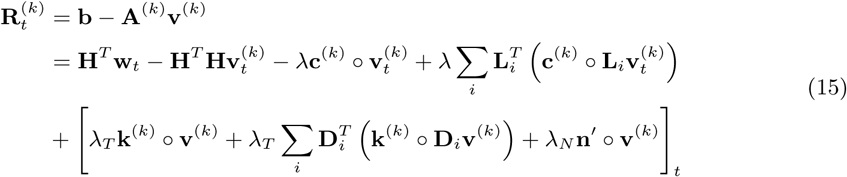

with **c** and **k** given by eqs. (5) and (6) and *λ*_*N*_ = 100*λ*. Another definition is

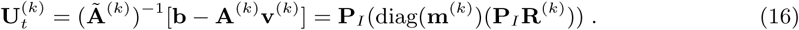

To evaluate the “goodness” of the iteration result, the following measure is introduced

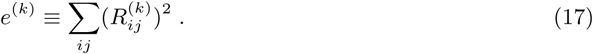

The resulting deconvolution algorithm is given in algorithm 1.

#### Algorithm 1

Deconvolution

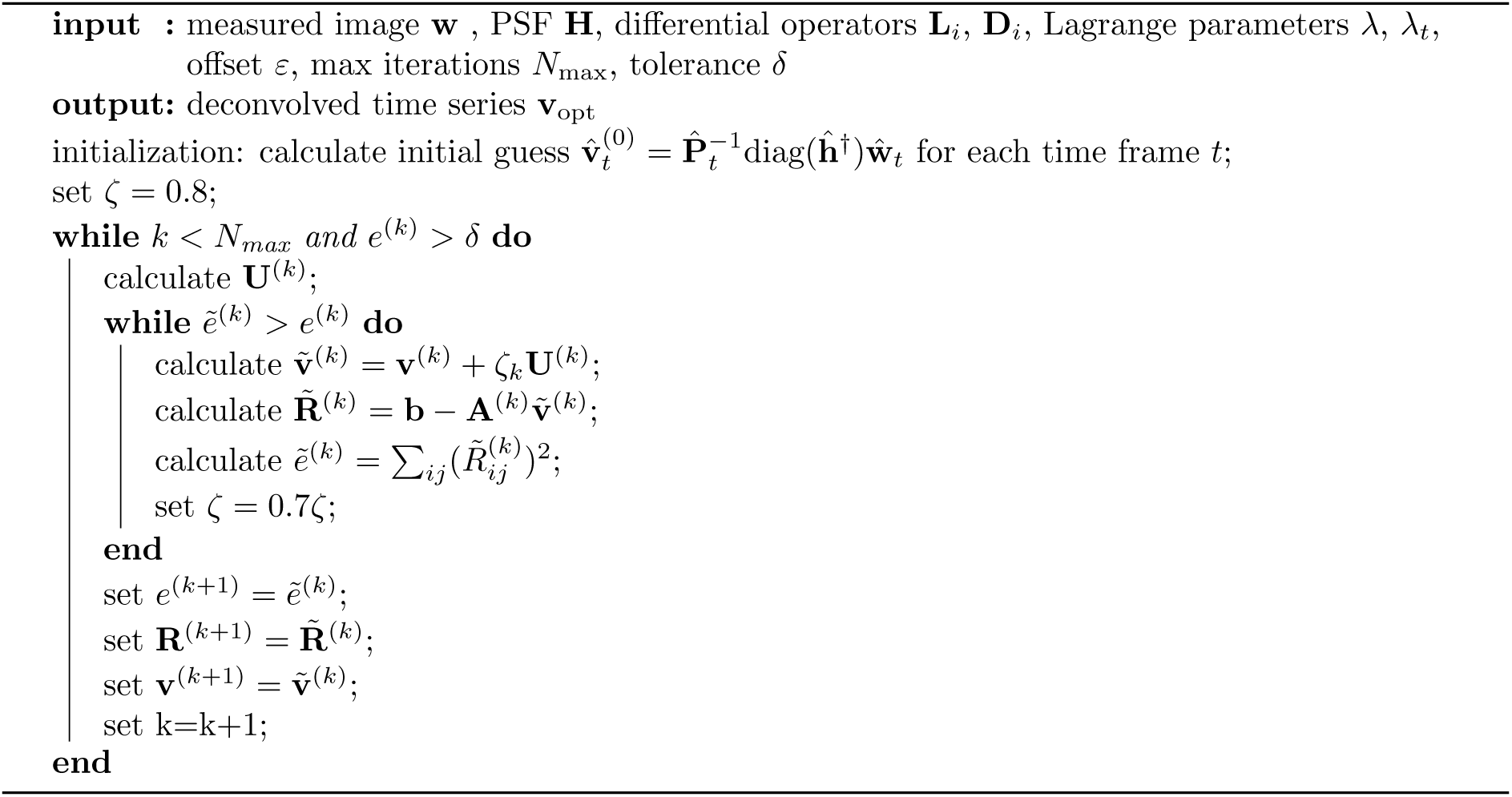

### A.2 Practical Notes

It should be noted that to facilitate notation, especially for the time dependence, matrix notation has been used throughout the previous section(s). However, it is computationally highly impractical to implement the algorithm exactly this way due to the enormous matrix sizes. In our implementation, the matrix product between a circular matrix and a vector, such as the image degradation denoted by **Hv** is actually implemented as a convolution *h* ∗ *v*, where *h*(*x, y*) is the point spread function and *v*(*x, y*) the image function depending on pixel values *x, y*. A matrix product with a transposed matrix such as **H**^*T*^ **v** is equivalent to convolution with a shifted kernel, or a cross correlation, i.e. *h* ⋆ *v*.

The parameters *λ, λ*_*T*_, *λ*_*N*_, *ϵ* as well as the maximum number of iterations *N* need to be determined empirically and can be widely different dependent on the type of data. In practice, *λ*_*N*_ is set to 100 · *λ* and *ϵ* is simply a small number such as 0.001. The best maximum number of iterations is often only *N* = 1. This leaves *λ* and *λ*_*T*_ to be chosen, which often work best when they are of similar order of magnitude. Thus, effectively, only one parameter has to be chosen.

The source code is written in Python and provided at `github/ipmi-icns-uke/XXX` (available after acceptance of manuscript).

## Notes

### Competing Interest Statement

The authors have declared no competing interest.

